# Infant Left Amygdala Volume Is Negatively Associated with Fecal Microbiota Diversity

**DOI:** 10.1101/2023.04.23.537273

**Authors:** Anna-Katariina Aatsinki, Jetro J. Tuulari, Eveliina Munukka, Leo Lahti, Anniina Keskitalo, Henna-Maria Kailanto, Saara Nolvi, Noora M. Scheinin, Jani Saunavaara, Riitta Parkkola, John D. Lewis, Niloofar Hashempour, Satu J. Shulist, Linnea Karlsson, Hasse Karlsson

## Abstract

**Introduction:** Rodent studies have addressed the importance of early life gut microbiota in the development of emotional and social functioning. Studies in human infants are still scarce, but associations with cognition and temperament have been reported. Neuroimaging studies have linked the amygdala with fecal microbiota diversity in infants, but crucially these studies have not covered the neonatal period, and the current study addressed this gap.

**Methods:** The study population included 65 infants drawn from the ongoing, general population based FinnBrain Birth Cohort Study. Brain MRI was performed around the age of one month (mean age 25 days). Fecal microbiota profiles (mean 68 days) were assessed by 16s rRNA amplicon sequencing at the age of 2.5 months.

**Results:** We found a negative association between infant left amygdala volume and alpha diversity (n=52, beta =-0.0043, p=0.034, adjusted for infant sex, breastfeeding, delivery mode, age during fecal sampling, age from conception during scan, and intracranial volume, Fig.1). Amygdala volumes were not associated with beta diversity (p=0.21), nor with the abundances of individual genera when adjusted for the same covariates and multiple testing.

**Conclusion:** Our results provide first evidence for associations between the brain and fecal microbiota in human neonates. Although the reported data do not allow investigation of underlying mechanisms, i.e. about the directionality of the hypothesized gut-brain connection, the reported connection encourages for future investigations of manifestations of gut-brain axis in early life.

## 1 Introduction

Gut microbiota modulation in early life in rodents is related to altered fear reactivity, social functioning, and anxiety-like behavior, that are phenotypes often linked to amygdala. For instance, neuronal activity of amygdala is altered in germ-free mice, i.e. mice grown free of microbes, indicating that early life microbiome contact is important for both amygdala functioning. The exact mechanism(s) mediating the above-mentioned connection is unknown, but the bidirectional communication between gut microbiota and brain have been shown to involve neural, metabolic and immune pathways (1,2). The presence of gut microbiome is crucial for normal neurodevelopment in rodents (3). Germ-free rodents, i.e. rodents living in microbe-free environment, show altered fear reactivity, neuronal activity of the amygdala (4) and impaired fear recall (5). Later colonization of the germ-free animals only partially reverses or does not reverse these phenotypes, suggesting that early life is an important time window for the bidirectional signalling within the gut brain axis and subsequent emergence of amygdala-dependent fear recall (4,5). The proposed pathways of microbiota-gut-brain axis communication include the vagus nerve, immune system functioning and metabolism of tryptophan (6).

The amygdalae are important brain regions for processing emotionally salient information also in humans. Gut microbiota alterations have been linked with temperament, behavioural problems and differences in social and communication skills in human infants (7–9). Moreover, in one-year-old infants, higher microbiota alpha diversity at age 1 year, i.e. the intra-individual diversity of bacterial species, was associated with larger volume of left precentral and left cingulate gyri, and left amygdala and poorer cognitive performance at the age of two years (10). Another study using the same sample reported reduced functional connectivity between the left amygdala and thalamic areas as well as between the anterior cingulate cortex and anterior insula. (11). Further, alpha diversity was found to associate positively with functional connectivity between the supplementary motor area and the inferior parietal lobule, and this connectivity also associated with cognitive outcomes (11). Collectively, prior reports imply that there are relevant links between the gut microbiome diversity, brain structure and function, and child phenotypes already during early infancy.

Prior studies provide clear indications that the bidirectional signalling between the gut microbiome and brain exist in humans, but they do not provide evidence of causality and are unable to confirm the directionality of the communication – from the gut to the brain, vice versa or both. Infant’s fecal microbiota composition is characterized by the Bifidobacterium and Bacteroides species, and is affected by multiple factors, including the mode of delivery, breastfeeding, exposure to antibiotics, and environmental factors (12). Importantly, the microbiota strongly diversifies during early life and especially after the cessation of breastfeeding (12), and thus the findings from later time points cannot be translated to early infancy in a straightforward manner. It also remains unknown how early on during development the associations between brain structure and gut microbiome diversity emerge.

In the current study, we tested whether neonatal amygdala volume associate with early fecal microbiota diversity and composition measured at 2.5 months. The amygdala is an early-developing structure that is central for fear processing, and has been implicated to associate with microbiota diversity in prior work with older infants (10). We hypothesized to find a positive association between amygdala volumes and alpha diversity (10).

## 2 Methods

The study was conducted according to the Declaration of Helsinki and was approved by the Ethics Committee of the Hospital District of Southwest Finland. All mothers provided written informed consent on behalf of their children.

To study the associations between infant microbiota and amygdala volumes, neuroimaging and fecal samples were obtained from healthy infants (n=56, boys =24, 43%) participating in the FinnBrain Birth Cohort Study (13). MRI scans were performed around the age of one month (age at scan: 25±7 M±SD days, estimated age from conception: 304±7 M±SD days, Table 1) and fecal samples were obtained around 2.5 months of age (68±14 M±SD days, Table 1). Key infant and maternal variables were obtained from Finnish Medical Birth Register (https://thl.fi/en/web/thlfi-en/statistics-and-data/data-and-services/register-descriptions/newborns). Breastfeeding was measures by questionnaires and was categorized as exclusive breastfeeding, partial breastfeeding, no breastfeeding, or breastfeeding ceased.

**Table 1.**
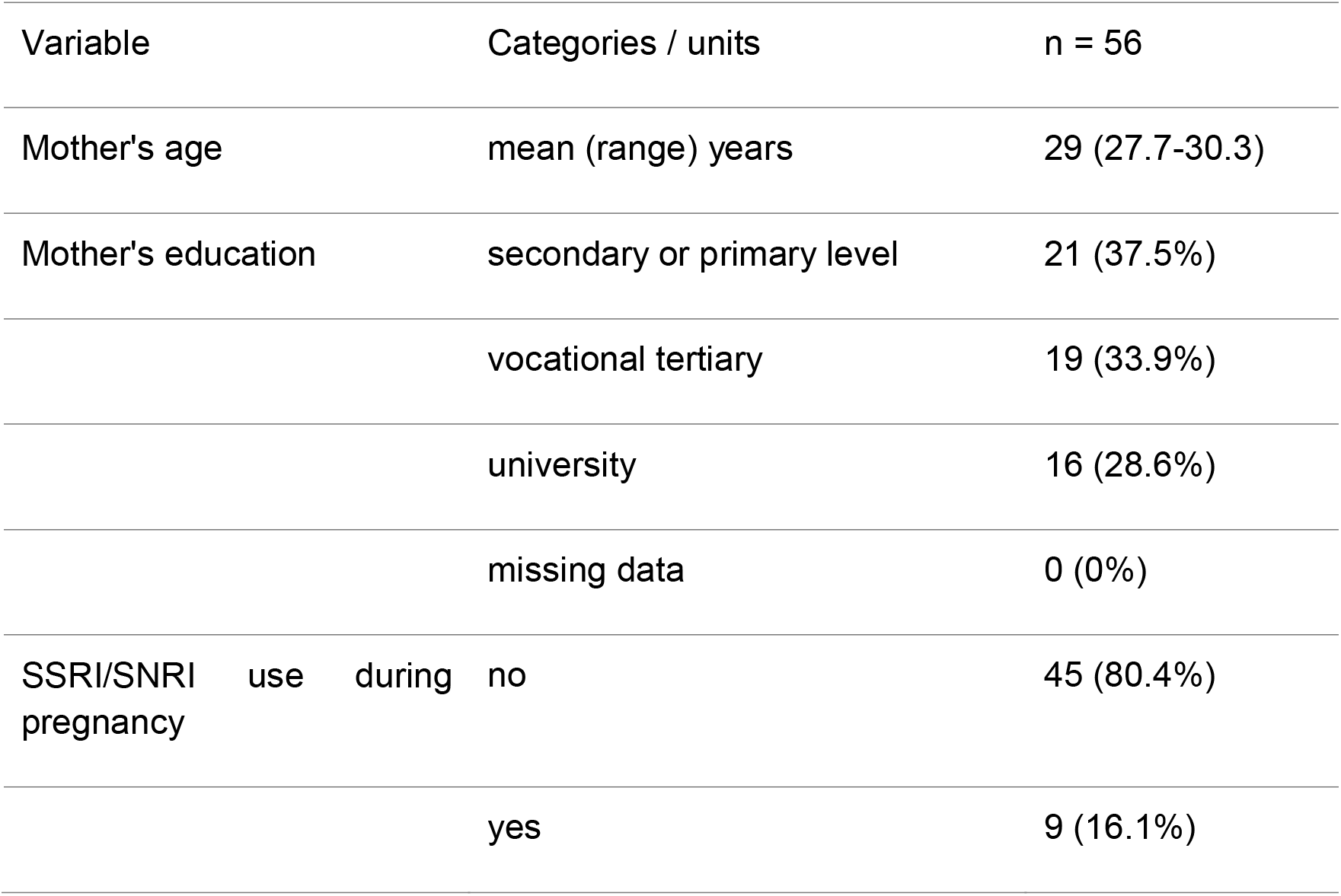
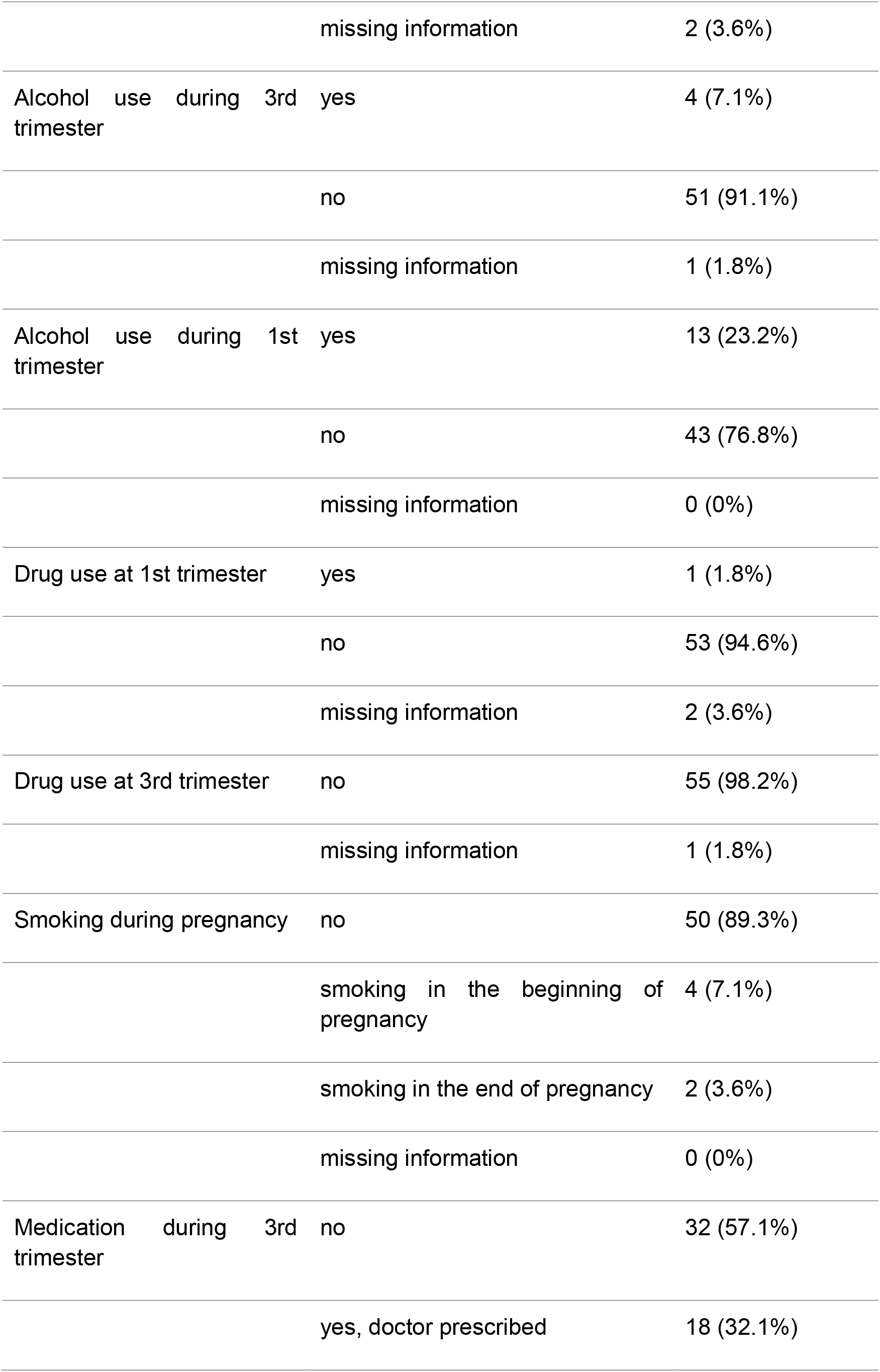
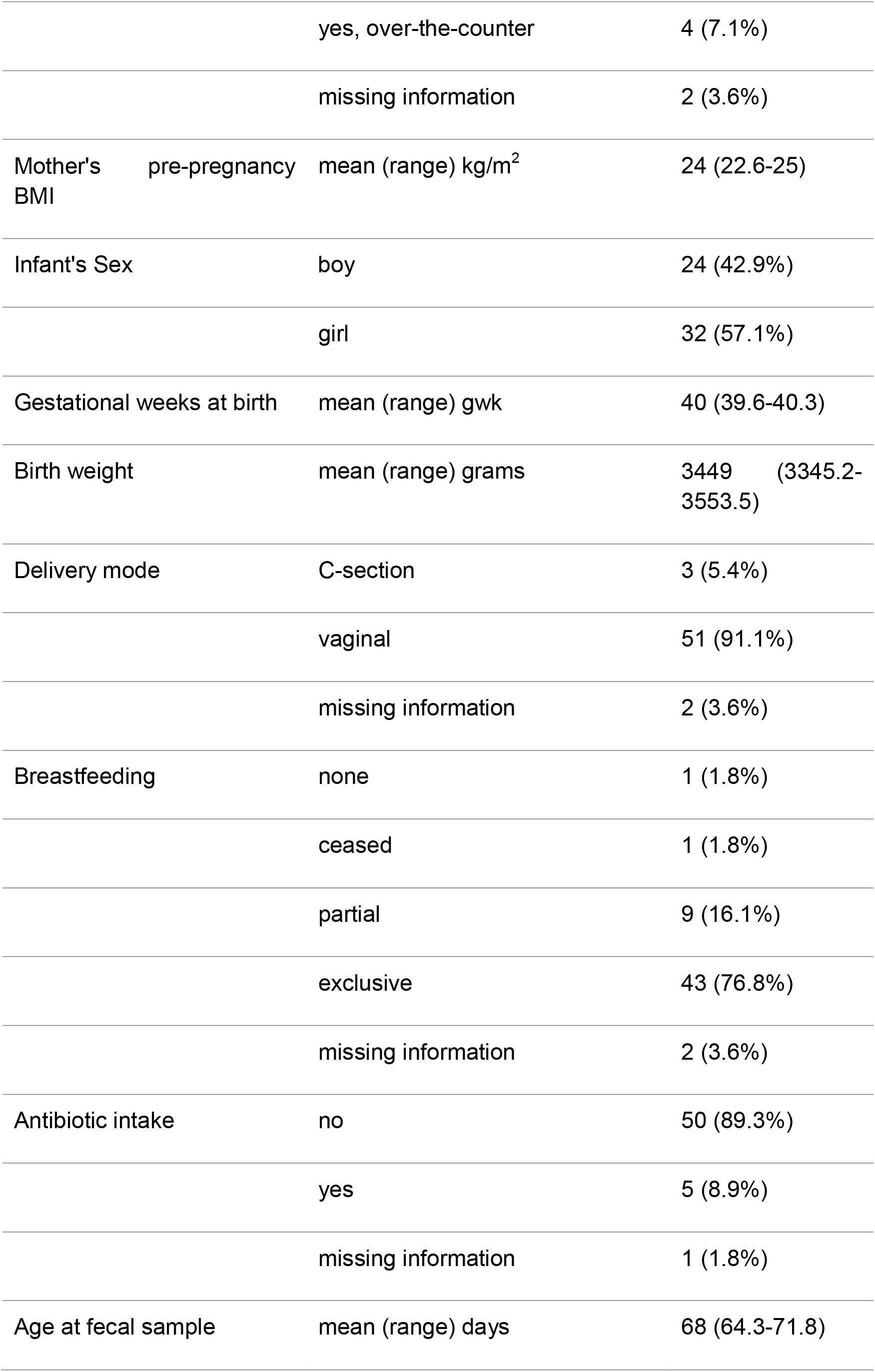
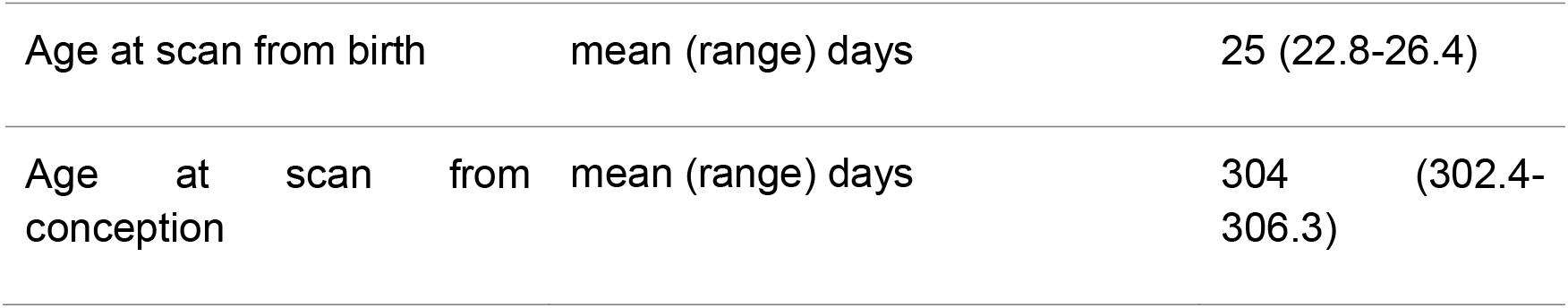
Description of maternal and infant characteristics.

MR imaging was performed during natural sleep with a Siemens Magnetom Verio 3T MRI scanner (Siemens Healthineers, Erlangen, Germany). The segmentation procedures have been described in our prior article (14). We used a previously published protocol for segmenting the bilateral amygdalae (15) and intracranial volume. Segmentation for each subject was done using a label-fusion-based labeling technique based on Coupé et al. (2011) and further developed by Weier et al. (2014) and by Lewis et al. (2019) (16–18).

Fecal samples were collected by parents at the infant age of 2.5 months as previously reported (19). To analyse the bacterial profiles of the fecal specimens, variable region V4 of the bacterial 16S rRNA gene was amplified and sequenced with an Illumina MiSeq system as previously reported (19). This sample had 210k reads on average (total 11771792, range 79200-940303). Shannon Index and number of observed OTUs were used to estimate intra-individual diversity and richness. Differences in genera abundances were analyzed with DESeq2 (20). Beta diversity, i.e. the variation in the overall community composition of the fecal microbiota across individuals, was tested with PERMANOVA using the Bray-Curtis dissimilarity with the adonis function from the vegan package (21).

Statistical analyses included regression models testing the association between bilateral amygdala volume and alpha diversity (Shannon Index). We first assured that both key variables were normally distributed. A priori chosen covariates of no interest included (infant sex, breastfeeding, delivery mode and age during fecal sampling, age from conception at MRI scan, and intracranial volume). The models were performed as follows:

Alpha / beta diversity / genera abundance ∼ Amygdala Left / Right + infant sex + age from conception at MRI scan + intracranial volume + breastfeeding + delivery mode + age at fecal sampling

Four subjects were missing some of the covariate data and were excluded from the analyses resulting in a sample size of n=52 for the statistical models. To test the robustness of the findings, the models with statistical significance (defined as p-value < 0.05) were assessed excluding the non-breastfed and c-section born infants from the analyses (n=48). Further, leave-one-out cross-validation tests were performed for the linear models with statistical significance. As the gut microbiota has been suggested to have sex-specific effects on neurodevelopment (22), sex interactions were also investigated by including an interaction term amygdala volume * infant sex in the linear models.

## 3 Results

The mean age of the mothers was 29±5 (M±SD) years and mean pre-pregnancy body mass index was 24 kg/m^2^ (SD=4) (Table 1). Eighteen mothers (32%) had prescription medication during the 3^rd^ trimester and nine mothers reported SSRI/SNRI use during pregnancy (16%). Two mothers (4%) reported smoking and four mothers (7%) reported alcohol consumption at the end of the pregnancy; none reported illicit drug use. Mean gestational age was 40±1 weeks (M±SD) weeks, all infants were born normal weight (3449±385 grams±SD), three infants were delivered with caesarean section (5%), and five infants (9%) had received antibiotic treatment on the neonatal ward. Most infants were breastfed at the time of stool sampling (exclusive breastfeeding: 77%, mixed feeding: 16%, no breastfeeding: 3.5 %, missing information 3.5 %; Table 1).

We found a negative association between infant left amygdala volume and alpha diversity (n=52, beta =-0.0043, p=0.034, adjusted for infant sex, breastfeeding, delivery mode, age during fecal sampling, age from conception during scan, and intracranial volume, Figure 1). This finding was similar in the subpopulation of vaginally born breastfed infants (n=48, beta = -0.0049, p=0.017, when adjusting for the same covariates). No interaction by infant sex was observed (p>0.3). No associations were detected between right amygdala volume and diversity (p=0.95), or between richness and amygdala volumes (p>0.6) in similarly constructed linear models. A leave-one-out cross-validation test further indicated that the left, but not right amygdala volume significantly improved predictions on fecal microbiota diversity in leave-out samples. Amygdala volumes were not associated with beta diversity (p=0.21), nor with the abundances of individual genera when adjusted for the same covariates and multiple testing.

**Figure. 1.**
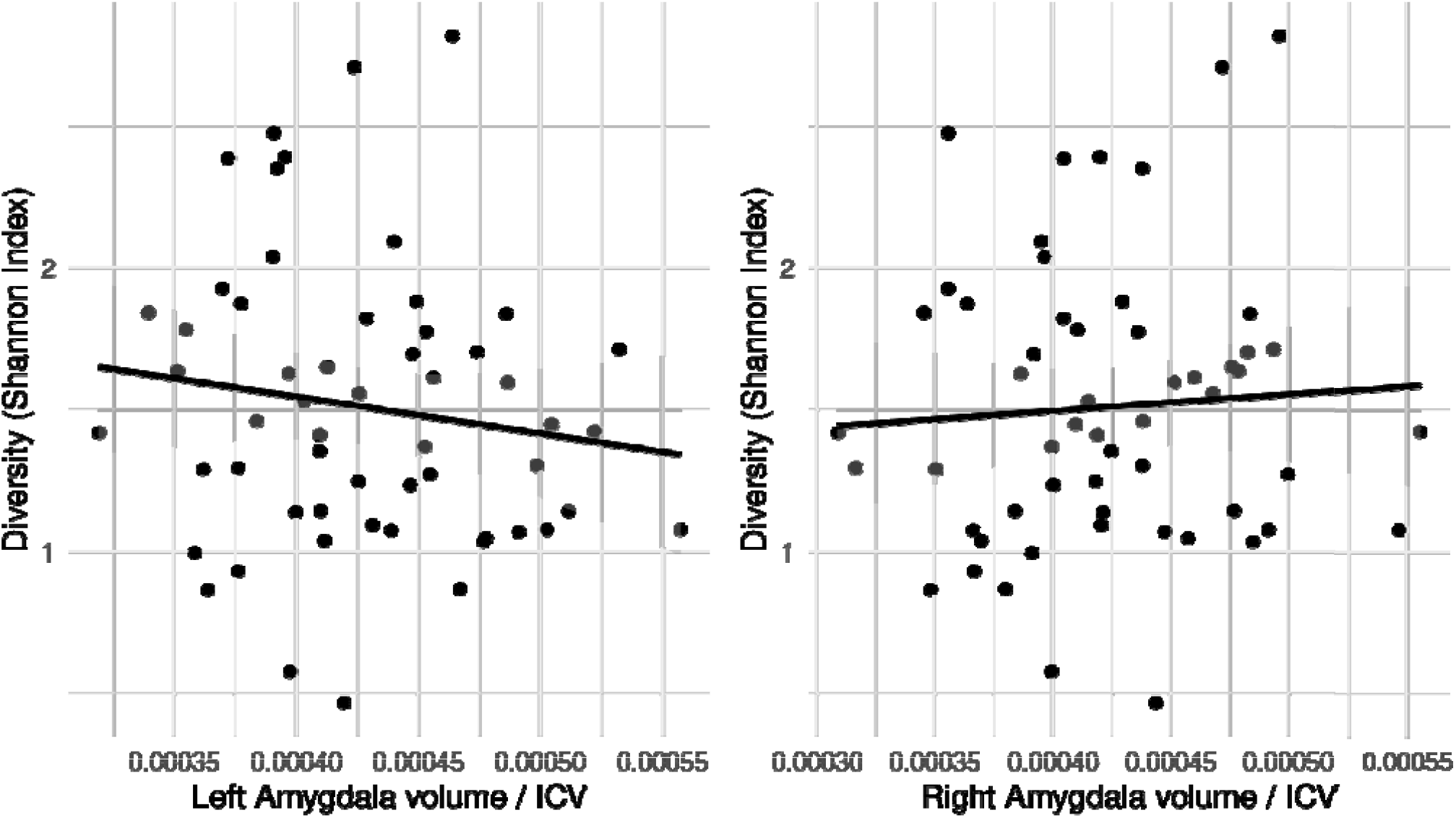
Gut microbiota alpha diversity associates negatively with left (A), but not with right (B) amygdala volume. The grey areas depict 95% confidence intervals.

## 4 Discussion

We found a negative association between infant left amygdala volume and alpha diversity, and no associations were found for right amygdala volume or between beta diversity and bilateral amygdalae. These results were contrary to our hypothesis that was based on prior work in older, 1-year-old infants (10), whose association was positive (albeit between alpha diversity at child age 1 year and brain volume at age 2 years). This implies that associations may be changing with increasing age but caution is warranted due to small sample sizes used, and larger scale replication studies are clearly needed.

It is well possible that the associations between amygdala and gut microbiome vary across the first year of life as both systems exhibit highly dynamic changes during this time (1,2). The potential connection between amygdala volume and microbiota may be modulated by various postnatal factors, such as breast milk composition (23) or immune challenge (24), which may underline the discrepancy between our results and the previous reports. Similarly to prior studies (10,11), our results implicate a unilateral association with left amygdala volume. Although the effect size between the volume of the amygdala and alpha diversity in both studies were small.

The significant association observed between alpha diversity and amygdala volume, and the simultaneous lack of association between beta diversity (community composition) and amygdala volume suggest that the associations between microbiota variation and amygdala volume relate to ecosystem-level features rather than taxonomic groups. Typically, later in development more diverse microbiota is more stable and potentially contributes to favorable health outcomes (25,26). However, contradicting this, increased fecal microbiota diversity has been reported to associate with worse cognitive development (10), and other studies investigating child cognitive or behavioral phenotypes and fecal microbiota characteristics usually report associations with fecal microbiota composition rather than with alpha diversity (8,9,27), which severely complicates the interpretation of the results across different age groups.

There are numerous factors potentially underlying the observed connections between brain development and gut microbiome. For instance, (endogenous) glucocorticoid exposure during pregnancy, that has been associated with both reduced amygdala volume (28) and the fecal microbiota composition of the progeny (19) should be investigated. Moreover, hypothalamus-pituitary-adrenal axis functioning, serotonin metabolism, and vagus nerve signaling from the gastro-intestinal tract may play a part in mediating the potential bi-directional microbiota-gut-brain axis signalling, among other factors (6). Future studies may benefit from complementing their designs with genetic analyses (14), additional biological measures (23,24), and inclusion of early exposures into their design (29). Longitudinal sampling, obtaining imaging and faecal samples at the exact same age, as well as larger sample size would potentially facilitate detection of biologically meaningful associations between amygdala volumes and fecal microbiome.

This is the first study on brain structure and gut microbiota in this age group, but the conclusions are limited by the modest sample size and a cross sectional design. Although gut microbiota diversity increases during infancy, it is significantly driven by the cessation of breastfeeding (12). Most of the infants in the present study were breastfed during imaging and fecal sample collection, which partially standardises the setting and is typical for young infants, but may also decrease the variation related to fecal microbiota diversity.

In conclusion, our results provide first evidence for an association between amygdala volume and gut microbiome diversity in human neonates. These results, together with prior studies in older age groups, encourage further investigation in longitudinal settings that would enable insights into underlying mechanisms, directionality of the hypothesized gut-brain connection, and later behavioural outcomes.

## 6 Author Contributions

LK, HK, AA, JJT designed the research. AA, JJT, AK, EM, SJL, HMK, NMS, JS, RP, NH collected the data. JDL designed and developed the MRI protocol and data processing tools. JS, JJT, RP, NH participated in the MRI data processing. EM, AK participated in fecal sample processing. AA, LL analysed the data. AA, JJT wrote the initial draft of the manuscript. AA, JJT, AK, EM, SJL, HMK, NMS, JS, RP, NH, SN, HK, LK, JDL critically revised the paper. All authors accepted the final manuscript.

## 8 Acknowledgments

We want to thank all the participating families and the FinnBrain staff and assisting personnel including Heidi Isokääntä, Minna Lamppu, and Katri Kylä-Mattila. We acknowledge Juha Pursiheimo for the collaboration regarding the 16S rRNA gene sequencing and Sami Pietilä for processing the raw sequences with QIIME pipeline.

## 9 Data Availability Statement

Due to Finnish federal legislation, the research data cannot be made available online, but data can potentially be shared with a Material Transfer Agreement as a part of research collaboration. Research collaboration requests can be directed to the Board of FinnBrain Birth Cohort Study. Please contact Linnea Karlsson.

